# UNWANTED AXON GROWTH: PTEN AND THE SUPPRESSION OF AXON PLASTICITY IN ADULT NERVES

**DOI:** 10.1101/2025.06.04.657750

**Authors:** Shane Eaton, Prashanth Komirishetty, Matt Larouche, Aparna Areti, Honyi Ong, Jose Martinez, Douglas W. Zochodne

## Abstract

In adults, peripheral nerves comprise bundles of disseminated motor, sensory and autonomic axons that are considered stable neuroanatomical units. The only exceptions to this established wiring are very distal terminal branches in target organs, such as skin. Here we provide a remarkable deviation from this state of affairs in the peripheral nerves of mice with a conditional knockout of sensory neuron PTEN (phosphatase and tensin homolog deleted on chromosome ten). PTEN is normally expressed in adult sensory neurons, particularly small IB4 nonpeptidergic subtypes and its knockdown after injury or during experimental diabetes improves axon regrowth. We studied Advillin Cre;PTEN null mice lacking PTEN in their sensory neurons. As might be expected, their harvested and cultured DRG neurons displayed enhanced neurite outgrowth *in vitro. In vivo*, these mice were healthy and had a normal sensory behavioural phenotype. However, the nerves of mice lacking sensory neuron PTEN were highly abnormal, with augmented clusters of small myelinated and unmyelinated axons populating endoneurial fascicles of their peripheral nerve trunks. The axon clusters did not disrupt normal fascicular anatomy but invested the epidermis with greater axon numbers. Within endoneurial fascicles, supernumerary axons formed regenerative units and expressed ongoing growth markers, unlike normal adult axons. This was not accompanied by rises in dorsal root ganglia (DRG) neuron numbers, indicating enhanced distal sprouting from parent neurons. Additionally, sprouting axons were electrophysiologically intact, generating rises in the amplitudes of sensory nerve action potentials. Despite this extensive regenerative activity of intact nerves, regeneration indices after superimposed injury were only modestly enhanced or unchanged. This unusual behaviour of adult sensory axons lacking a single growth-suppressive molecule may identify insights into what molecular constraints the nervous system normally utilizes to suppress inappropriate plasticity.

---

Structural stability of the nervous system may be essential to retain the fidelity of connections and to avoid inappropriate or misdirected connections. Neurons have long been classified as either within a stable, neurotransmitter role or, after axonal injury, a growth mode expressing regeneration associated genes (RAGs). In the CNS (central nervous system) development and learning likely involve structural remodeling, as does recovery from injury. In the PNS (peripheral nervous system) most depictions characterize a fixed ‘wiring’ structure that connects the periphery to the CNS. Inappropriate axon growth behaviour or misdirection might generate ectopic discharges or neuropathic pain. Recently, however, an exception to this state of affairs has been recognized that involves very distal, terminal sensory axons that innervate the epidermis ^2,4^. These axons ramify between keratinocytes in highly irregular, but spaced trajectories that suggest ongoing plasticity and growth. Such behaviour might be expected given their residence within an endorgan that routinely migrates keratinocytes to superficial skin layers then sheds them. Ongoing axon plasticity is likely essential to keep apace with normal keratinocyte loss. Axons in the epidermis of mice and humans express markers of growth and regeneration, not normally expressed in ‘stable’ axons within the parent nerve trunk ^2^. In more proximal nerve trunks ongoing growth of axons is not a feature.

PTEN (phosphatase and tensin homolog deleted on chromosome 10) is a well known tumour suppressor phosphatase molecule that dephosphorylates PIP3 (phosphatidylinositol (3,4,5)-triphosphate) to PIP2(phosphatidyl inositol (4,5) on the membranes of cells to inhibit PI3K/pAkt (phosphoinositide 3-kinase/protein kinase-B) growth signaling ^30^. Mutations downregulating PTEN are associated with cancer susceptibility. Moreover, PTEN is expressed physiologically in sensory neurons, with heightened expression among small caliber IB4 nonpeptidergic subtypes ^7^. PTEN is not among the classical ‘RAGs’ but declines in pPTEN (phosphorylated PTEN) following axotomy were observed in the nuclei of large and medium sized neurons ^7^. However, the expression of PTEN in small caliber nonpeptidergic neurons was remarkable and correlated with their known properties of relative growth impairment ^12,28^. The coincidence of slower intrinsic growth and higher PTEN expression suggests an ongoing role for PTEN in dictating growth behaviour in DRG sensory neurons. Exogenous knockdown of PTEN in wild type mice enhances neurite outgrowth of adult sensory neurons and improves indices of regeneration *in vivo* ^7^. These findings correlate with those identified in optic nerve and corticospinal tract regeneration of the CNS ^13,20,21^. PTEN is degraded in neurons by NEDD4, a constitutively expressed ubiquitin E3 ligase and NEDD4 knockdown, by preserving PTEN levels, impairs outgrowth in sensory neurons ^6^. Further, in mice with chronic experimental diabetes that develop sensory polyneuropathy resembling that of humans, PTEN expression is upregulated. PTEN knockdown in diabetic models improves injury-induced regeneration. This is important because superimposed focal injuries are common in diabetes and diabetic nerves have an added regenerative deficit. PTEN inhibition or knockdown, however, also reverses the degenerative retraction of epidermal axons that characterizes polyneuropathy in diabetic animals ^22,26^. Overall, these roles in experimental diabetes provide yet another link between PTEN expression and regenerative impairment.

Here we characterized a mouse model of selective PTEN deletion, Advillin Cre;PTENflx/flx in sensory neurons. We noted an unexpected phenotype of intra-endoneurial axon sprouting with an apparent normal sensory behavioural phenotype. Further, there was remarkable uneven expansion of myelinated and unmyelinated axon populations within intact nerves without overall nerve distortion. These expansions comprised sprouts from existing neurons, not accompanied by additional DRG sensory perikaryal numbers. The axon clusters differed from Remak bundles but were typical of regenerative units ultrastructurally. Their features were variably myelinated axons, expression of regeneration markers and the expanded populations were electrically excitable. Invasion beyond the endoneurium was not identified but there was enhanced epidermal innervation of the hindpaw footpads. The findings suggest a unique pattern of axon sprouting in otherwise intact peripheral nerve trunks linked to the deletion of a single growth suppressive molecule.

## Results

### Model characterization

Conditional sensory neuron PTEN-/-mice were bred from crossing mice expressing Advillin linked Cre recombinase and PTEN flx/flx mice (AdvCre;PTENflx/flx). The offspring expressing both transgenes lacked PTEN expression in DRG sensory neurons (**Figure 1A-C**). Mice did not exhibit abnormalities in feeding, weight gain or general activity. This mouse line was developed independently by our laboratory at two University sites over an approximate 15 year timespan. In our first cohort, we noted loss of the original conditional PTEN knockout phenotype during breeding despite initial characterization. At the new site, an independent model cohort was re-derived from new lines, yielding an identical mouse phenotype. Analysis of mice with loss of the PTEN deletion genotype from the original first cohort breeding line also provided a second control cohort of animals (**Supplemental Figure 1**). The rederivation of the line allowed us to confirm an identical phenotype from two separate breeding protocols.

**Figure 1:**
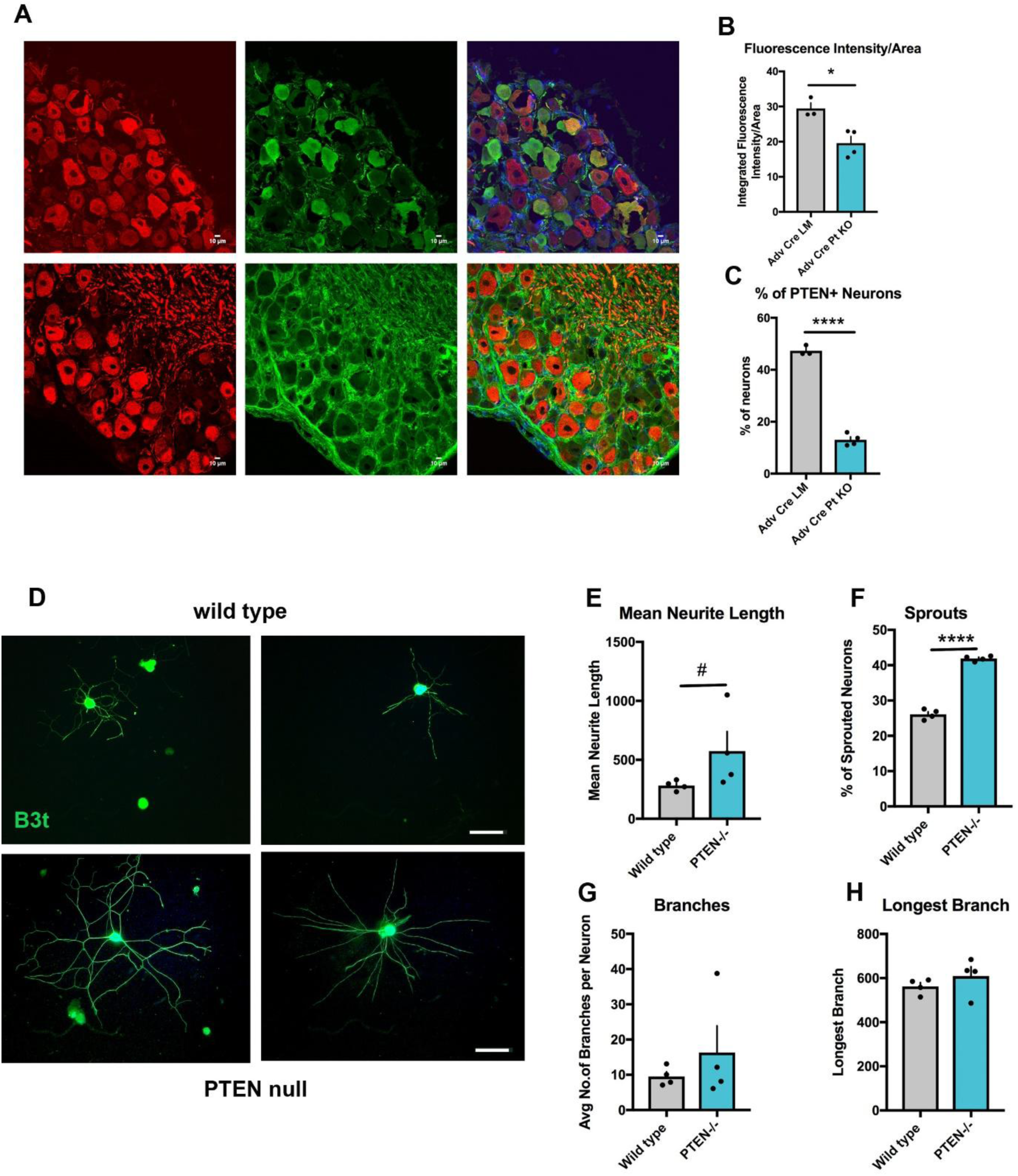
PTEN is deleted in DRG sensory neurons that display heighted outgrowth and sprouting. (**A**) Transverse sections of lumbar DRGs in wild type (top row) and sensory neuron PTEN-/-mice (bottom row) stained with antibodies directed against neurofilament (red) and PTEN (green) with merge. Note the loss of neuronal PTEN staining in the PTEN-/-mice [Bar=10 µm]. (**B,C**) Quantitation of immunohistochemical staining of DRGS showing the loss of fluorescence in staining for PTEN (B) and the decline in profile counts of the proportion of PTEN expressing neurons (**C**). (**D**) Examples of primary sensory neurons *in vitro* stained with βIII tubulin harvested from wild type mice or mice with sensory neuron PTEN deletion. (**E-H**) Quantitative neurite outgrowth from adult lumbar DRG neurons from wild type and sensory neuron PTEN-/-mice. PTEN deletion was associated with rises in neurite length and sprouts. [(B) *p=0.014; (C) **** p<0.0001, two tailed Student’s t-tests, n=3 wild type, 4 PTEN-/-; (E) # p=0.028, one tailed Mann Whitney, n=4/group; (F) ****p<0.0001 two-tailed Student’s t-test, n=4/group].

Harvested naive DRG neurons *in vitro* from mice without a preceding nerve injury had growth properties that were compared to neurons from PTEN-/-mice. The latter exhibited higher mean neurite outgrowth and sprouting. Longest branches were comparable between the two groups (**Figures 1D-H**).

Taken together the findings confirmed loss of PTEN expression in the model and supported previous work identifying heightened properties of neurite outgrowth in sensory neurons with PTEN inhibition or knockdown.

### Nerve structure, myelinated axon morphology, ultrastructure

The most remarkable features of the sensory neuron PTEN-/-mice were the histological characteristics of their peripheral nerves. In both the sciatic nerve and its distal sensory branch, the sural nerve, the overall fascicular area was greater in mice lacking PTEN-/-(**Figure 2A-C**). This rise in area was accounted for by rises in myelinated axon numbers while retaining near normal axon density. On inspection of transverse sections of these nerves, in addition to enlargement, we noted discrete enlarged zones of smaller myelinated axons separating the larger myelinated axon pools into sectors. These zones contained not only smaller myelinated axons but unmyelinated axons, confirmed below by electron microscopy (EM). A myelinated axon size histogram showed rises in the proportions of the smallest categories of axons in PTEN-/-mice but also increases in the proportion of larger myelinated axons.

**Figure 2:**
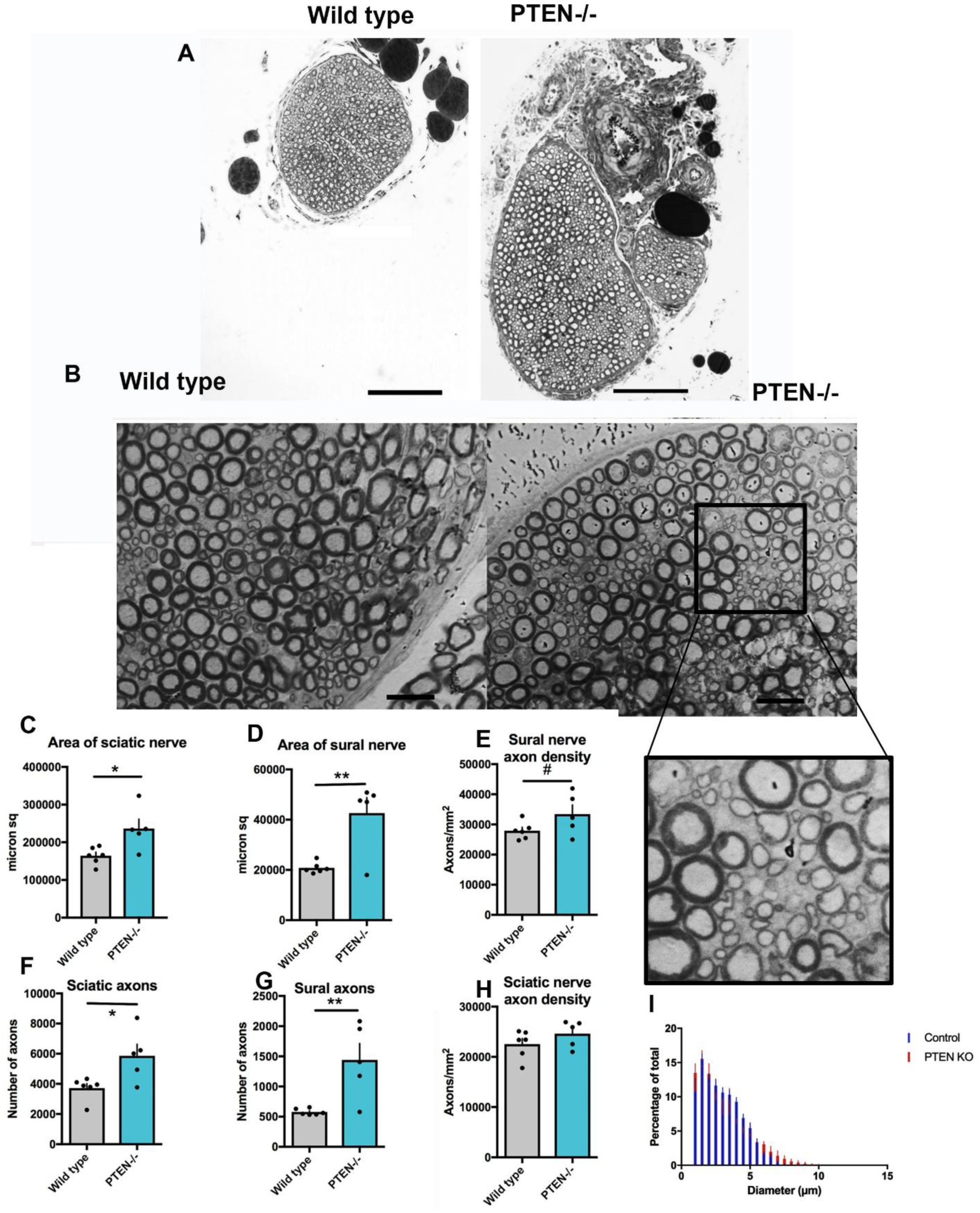
PTEN deletion in sensory neurons is associated with nerve enlargement, increased axon investment and clusters of small caliber axons. (**A,B**) Semithin LM images of sural nerves at lower and higher magnification showing enlargement of the nerve in PTEN-/-mice with supernumerary clusters of small axons (inset). [Bar=200 µm for A, 20 µm for B]. (**C-H**) Quantitation of sciatic and sural nerve areas, numbers of axons and axon density. Sciatic and sural nerves were larger in area in the PTEN -/- mice. (**I**) Myelinated axon diameter histogram of wild type (blue) superimposed with PTEN-/-data (red). The most marked rise in myelinated axon numbers was in the smallest size category (left). [(C) *p=0.018; (D)**p=0.004; (F) *p=0.02;(G)**p=0.007 two-tailed Student’s t-tests; n=6 wild type, 5 PTEN-/-; (E)#p=0.05 one-tailed Student’s t-test, n=6 wild type, 5 PTEN-/-].

Ultrastructural EM studies confirmed that the contents of axon pools separating larger myelinated fibers consisted of a variety of small axon collections (**Figure 3**). Some were small nascently myelinated axons indicative of an early myelinated sprouts. These were most often singly distributed but adjacent to single or groups of unmyelinated axons. Among distinguishing features of these clusters from normal wild type Remak bundles was their close axon adherence to one another without discrete separation by leaves of Schwann cell (SC) cytoplasm. These resembled arrangements classically described as regeneration units by Morris et al ^15-18^. Overall, the absence of SC separation of fibers, variable numbers of axon profiles within them and accompanying singly distributed unmyelinated differentiated the profiles from Remak bundles of wild type mouse nerves.

**Figure 3:**
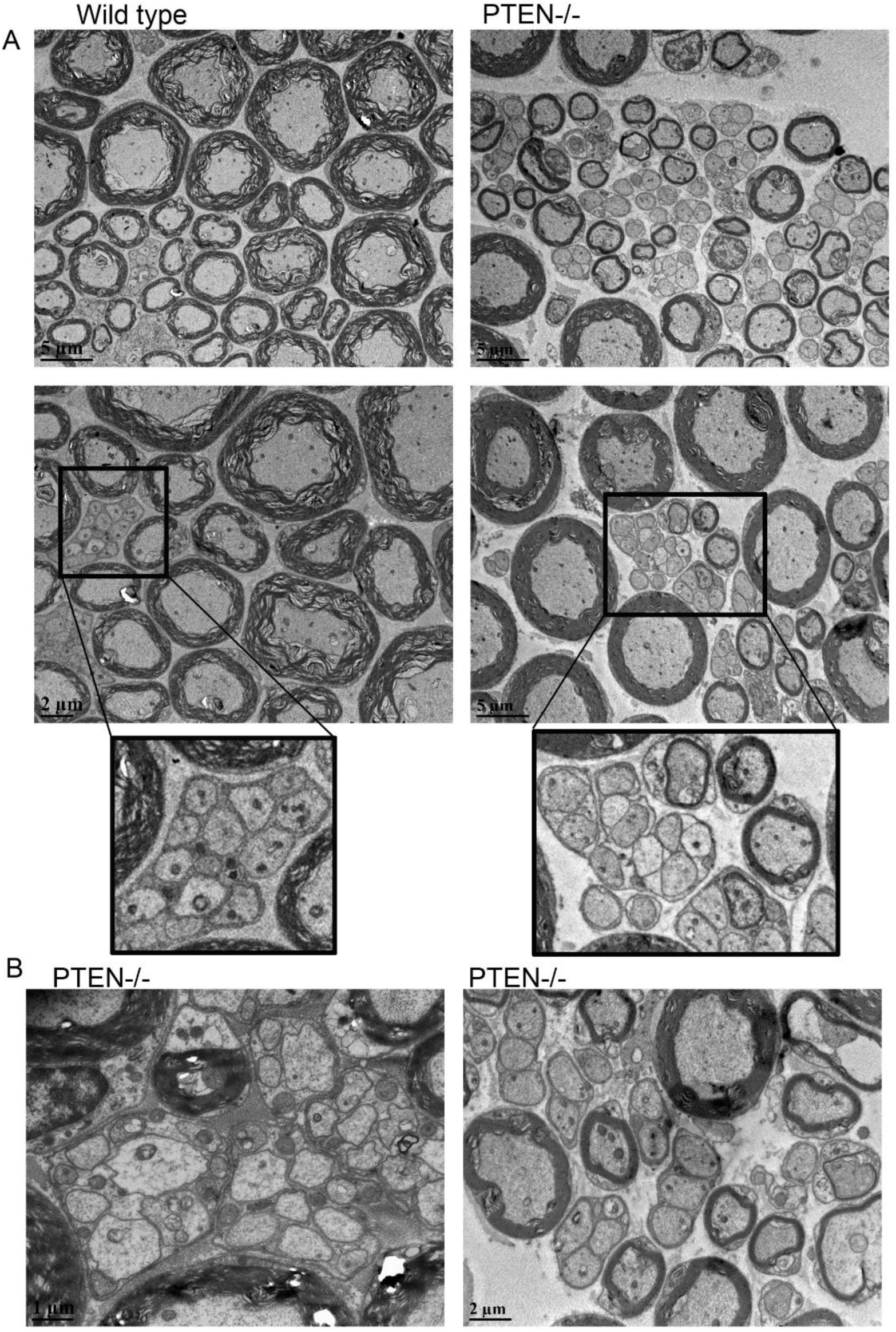
Adult sciatic nerves from sensory neuron PTEN-/-mice have zones of regenerative clustering. (**A**) Comparison on the EM ultrastructure of wild type and PTEN-/-mice at lower and higher magnification with insets. The PTEN-/-samples have clusters of small myelinated axons admixed among larger myelinated profiles. Unmyelinated axons are illustrated in common basement membrane groupings or as single profiles. The number and format of the clusters differ from normal orderly Remak unmyelinated bundles illustrated in the wild type nerves. (**B**) Additional high magnification images of regenerative cluster formation in PTEN-/-mouse nerves. Note the variation in axon maturity and myelination, a feature of regenerative clusters [Bars=1-5 µm].

Collectively these findings identified a remarkable intra-endoneurial myelinated and unmyelinated axon expansion in clusters that also involved more mature, larger axons. The findings suggested an ongoing level of axon plasticity of relatively long duration.

### Regenerative clusters of intact nerves

Immunohistochemical studies confirmed the above findings. We analyzed the association of myelination profiles, labelled with MBP, a myelin protein, with axon PGP 9.5 labelled profiles. Circular MBP profiles containing PGP 9.5 axons within their centers were prominent and correlated with the light microscopic profiles described above. Unlike wild type nerves however, nerves from PTEN-/-mice had distinct separated pools of PGP 9.5 axons with small MBP investments, or axons lacking myelin at all (**Figure 4A,B**). Quantitation confirmed a significant rise in the numbers of unmyelinated, PGP 9.5 staining without MBP, in the nerves of PTEN-/-mice (**Figure 4C**). While small groups of unmyelinated axons were expected and corresponded to normal Remak bundles in the wild type mice, these were more regularly spaced and much less variable in content.

**Figure 4:**
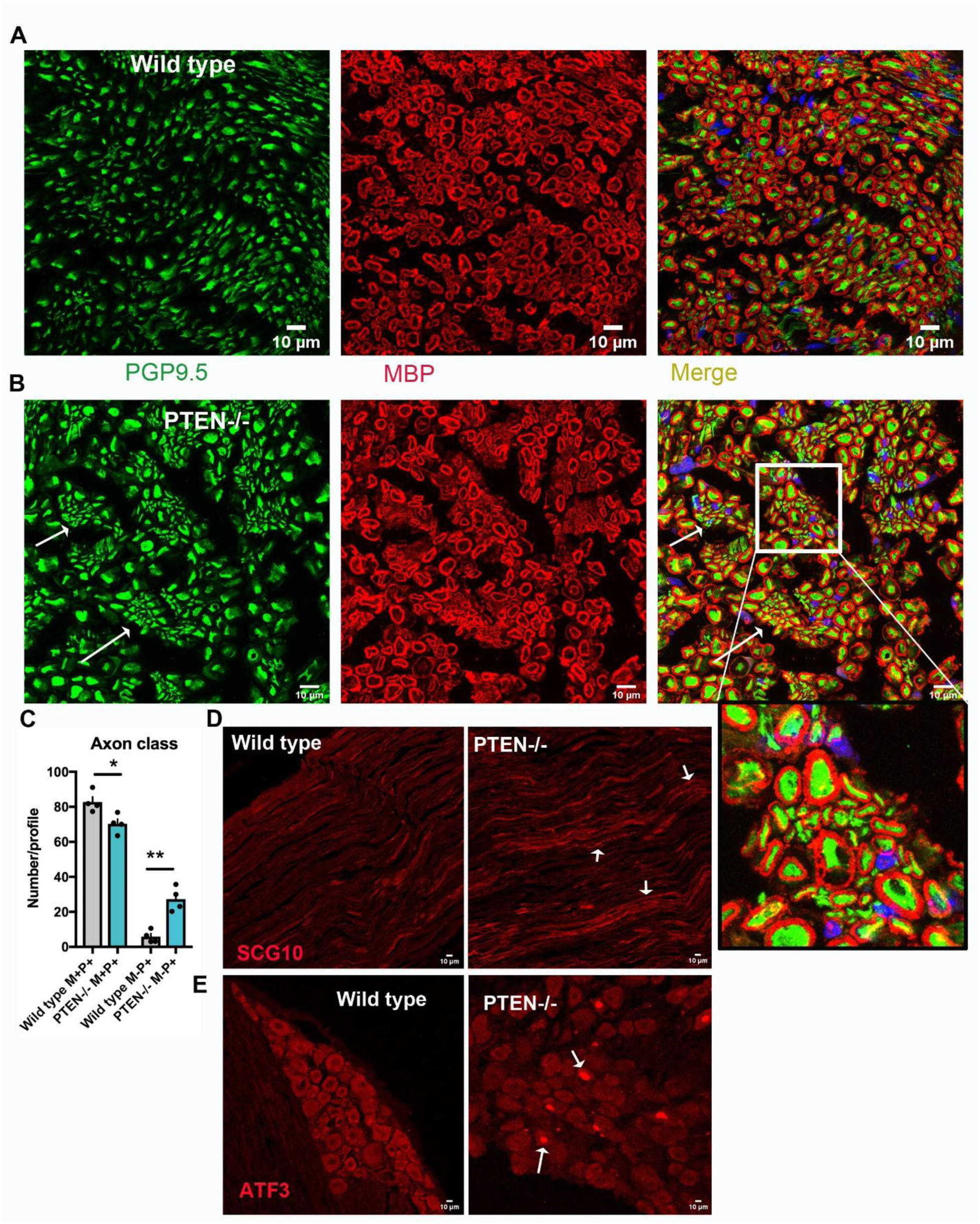
Clusters of small axons form adjacent to larger caliber myelinated axons. (**A**) Images of axon profiles (PGP9.5, green) colabelled with myelin (MBP, red) in wild type mice indicate regular and orderly spacing of myelinated fibers. Remak bundles are less prominent. (**B**) Axon profiles include several clusters of small caliber axons, some myelinated others without an MBP labeled myelin sheath. Enlarged inset highlights a regenerative cluster. (**C**) Quantitation of axon profiles with (M+P+) or without (M-P+) associated MBP labeled myelin sheaths. Sensory neuron PTEN-/-profiles with a myelin sheath were fewer in numbers (profile counts) whereas those without a myelin sheath were more frequent, indicating a rise in umyelinated axon numbers. (**D)** Longitudinal sections of intact nerves showing axons labeled with the regeneration marker SCG10 in PTEN-/-mice. Wild type mice had fewer, nonspecific profiles [Bars=10 µm]. (**E**) Sections of lumbar DRG stained with ATF3, a nuclear regeneration associated gene (RAG) protein. Nuclei in the PTEN-/-mice identified rises in nuclear ATF3 imunoreactivity, not seen in the wild type mice [Bars=10 µm]. [(C) *p=0.02; **p=0.0016, two tailed Student’s t-tests, n=4/group].

Nerves from PTEN-/-mice also exhibited axons stained with SCG10, a marker of sensory axon regenerative outgrowth (**Figure 4D**). SCG10 immunoreactivity was much less common in wild type nerves and usually nonspecific without discrete axon labelling. Finally, we stained parent lumbar DRG neurons with an antibody directed against ATF3, a marker of regenerative activation in sensory neurons routinely used to identify neurons having undergone axotomy injury. Despite being intact, PTEN-/-mice exhibited ATF3 nuclear staining indicating regenerative activation (**Figure 4E**).

Taken together these findings indicate a unique activation of status of intact nerves in the absence of PTEN, with ongoing axon sprouting, expression of an axon regenerative marker and evidence of regenerative reprogramming of their parent perikaryal.

### DRGs, epidermal innervation, excitability and axon function

Heightened sensory axon outgrowth in nerve trunks might be a result of enhanced numbers of parent sensory neurons in DRGs. Given the lack of evidence that normal DRG sensory neurons in adults are capable of expansion, this possibility seemed unlikely. However to address this possibility, we obtained profile counts of DRG neurons in lumbar DRGs from wild type and PTEN-/-mice. DRGs were labelled with βIII tubulin (B3T) as a marker for all sensory neurons or IB4 to label the nonpeptidergic populations. Neuron hypertrophy was not evident and counts of neurons with both labels were comparable between the mice (**Figures 5A,B**). Moreover the proportion of IB4 neurons compared to overall neuron numbers was similar between the wild type and PTEN-/-mice (**Figure 5B**). This is important because IB4 neurons normally have very prominent PTEN expression. These findings supported the concept that supernumerary axon numbers arose from sprouting, not from new neurons.

**Figure 5:**
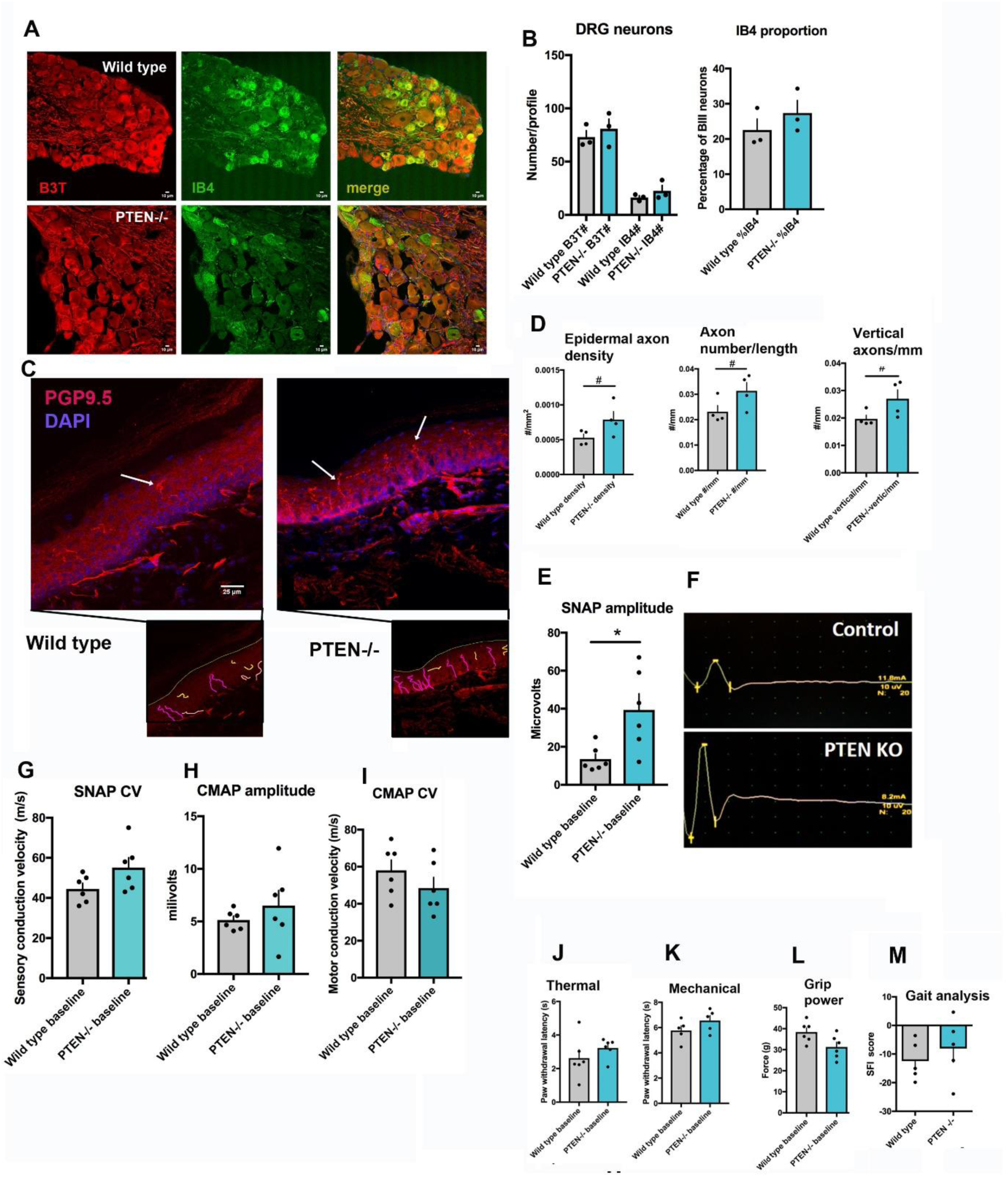
DRGs have preserved neuron numbers and proportions in sensory neuron PTEN -/- mice, have normal behaviour measures of nerve function but have increased epidermal axon density and higher sensory nerve action potential amplitudes (SNAPs). (A) Images of lumbar adult DRGs immunostained for βIII tubulin (B3T, red) and colabelled with IB4 (green). IB4 staining is more prominent in smaller caliber neurons. (B) Quantitation of the relative profile numbers of βIII tubulin neuron profiles and IB4 neurons indicating no difference in profile numbers or the proportion of IB4 neurons. (C) Images of hindpaw footpads and their epidermal innervation with axons labeled by PGP9.5 (examples shown by arrows)[Bar=25 µm]. The insets below illustrate axon tracing samples for quantitation. (**D**) Quantitation of epidermal innervation by density, numbers of axons per length and numbers of vertical axons per length (45-90^0^ angle from dermal-epidermal border). Sensory neuron PTEN-/-mice had rises in epidermal innervation. (**E-I**) Multifiber electrophysiological studies in wild type and sensory neuron PTEN-/-mice. Mice lacking PTEN had rises in the amplitudes of their sensory nerve action potentials (SNAPs), illustrated in (M), with comparable sensory conduction velocities, motor CMAP amplitudes and motor conduction velocities. (**J-M**) Behavioural functional measures comparing wild type and sensory neuron PTEN-/-mice. Hindpaw mechanical sensation, thermal sensitivity, grip power and gait analysis were comparable between the groups [(D) density #p=0.044, number/length #p=0.047, vertical #p=0.041, one-tailed Student’s t-tests, n=4/group; (E) p=0.017, two-tailed Student’s t-test, n=6/group]

Next we examined the sensory neuron tree at its distal reaches, epidermal skin innervation. At this site, unlike other parts of the PNS, some degree of plasticity and ongoing growth is not unexpected and previously reported in our group ^2,3^. Epidermal axon density, numbers per length and vertically directed axons were higher in number in the PTEN-/-mice than wild type controls (**Figures 5C,D**). These findings confirmed that distal outgrowing axon sprouts heightened innervation of the epidermis.

We queried whether the newly identified nerve populations of sensory axons were electrically excitable. Sensory nerve action potentials (SNAPs) recorded from digital stimulation and sciatic nerve recordings identified a substantial rise in their amplitude in PTEN-/-mice compared to wild type mice (**Figures 5E,F**). Sensory conduction velocities (CVs) were comparable as were the amplitudes and motor conduction velocities of the compound muscle action potentials (CMAPs) recorded in the same nerve distribution (**Figures 5G,H,I**). Given that SNAP amplitudes reflect numbers of electrically active myelinated sensory axons at the recording site, these findings match the morphological data indicating sensory axon expansion. They also indicate ongoing maturation of these sprouts since SNAPs are generated by myelinated, not unmyelinated axons. Since CV is measured to the fastest conducting larger caliber sensory axons that generate the SNAP, unchanged CVs indicate that new myelinated axons of yet larger caliber or faster CVs, were not a product of supernumerary axon growth. Constraints in growth normally limit the extent of axon radial growth, myelination and nodal properties and these persist in the setting of PTEN deletion. The findings also confirmed unchanged motor axon numbers and characteristics.

Finally we tested whether the expansion of sensory axons altered sensory or motor behaviour in mice. Wild type and PTEN-/-mice had comparable sensitivity to mechanical or thermal stimulation, comparable hindpaw grip power and similar normal sciatic functional index (SFI) metrics from gait analysis (**Figures 5J-M**). These findings indicated that ongoing sprouting of axons did not change the sensory behavioural phenotype. Given that sprouts arise from a fixed number of parent neurons, additional sensory input may not be generated by sprouting, a constraint of interest in these mice.

### Axon regeneration

In wild type mice, PTEN inhibition is associated with increased regeneration *in vivo* when PTEN inhibitors or knockdown are applied ^1,7^. To evaluate whether PTEN deletion in sensory neurons impacts regenerative indices we compared mouse behaviour, electrophysiology and axon numbers distal to a sciatic nerve crush after 14 and 28 days. As in previous work, nerve crush was associated with lower thresholds for mechanical and thermal sensation, a hyperalgesic response likely contributed to by collateral sensory axons ^5,19^. At both 14 and 28 days following crush, changes in both forms of sensation changed in parallel between the wild type and PTEN-/-mice (**Figure 6A,B**). Mechanical sensation was comparable at both time points. At 14 days, but not 28 days, thermal sensation was improved (less hyperalgesic) in PTEN-/-mice. Hindpaw grip strength and gait SFI measures similarly declined after injury with later recovery at 28 days but no difference was observed between the groups of mice (**Figure 6C,D**). SNAP and CMAP amplitudes and conduction velocities declined after crush with some recovery by 28 days with similar trajectories between the groups (**Figure 6E-H**).

**Figure 6:**
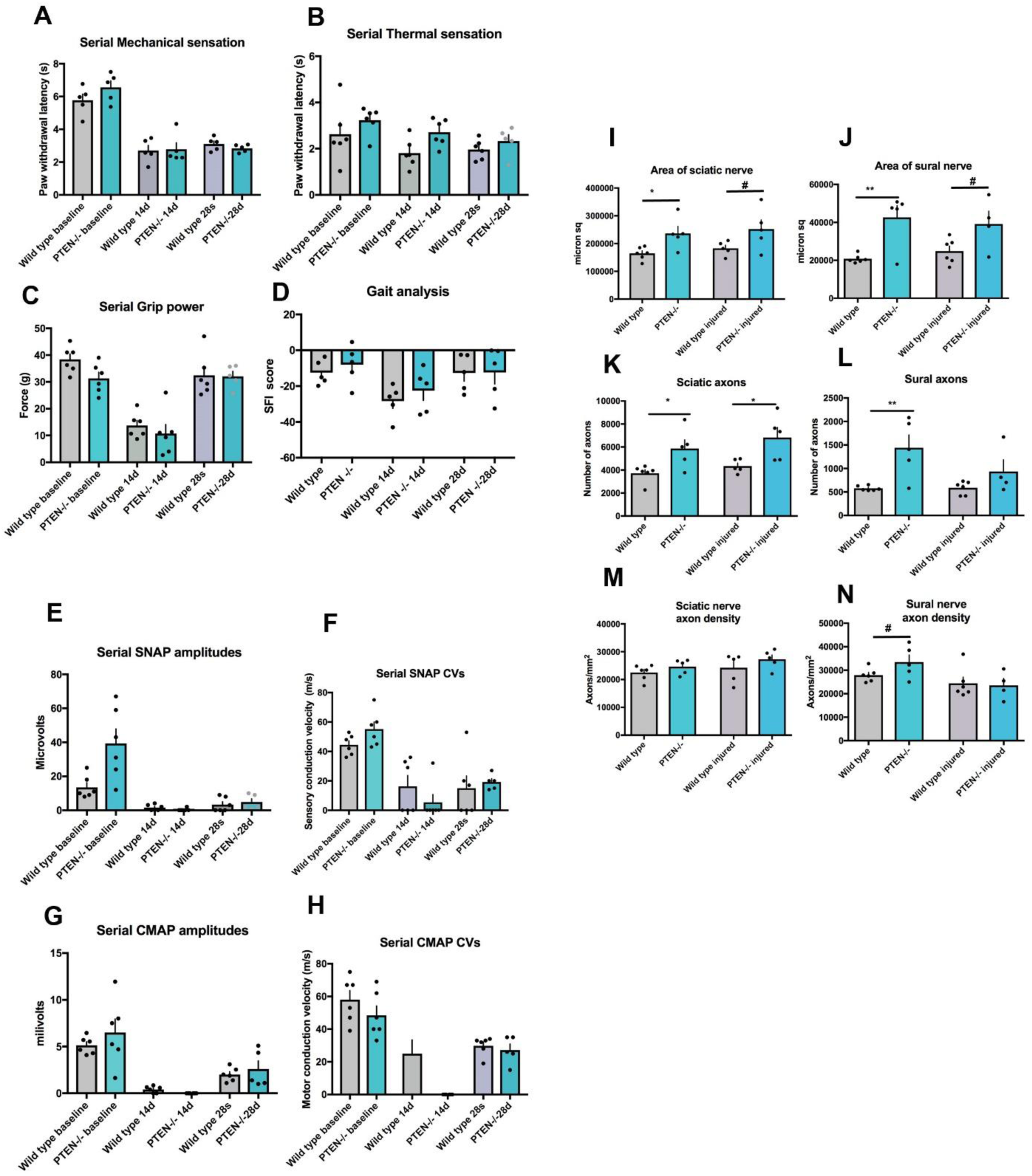
Mice with sensory neuron PTEN deletion have comparable recovery of sensory and motor abnormalities and electrophysiology after nerve injury but with persistent rises in sciatic myelinated axon numbers. (**A-D**) Serial measures of mechanical and thermal sensation, hindpaw grip power and gait analysis following sciatic nerve crush comparing baseline, 14d and 28d results. Wild type and sensory neuron PTEN-/-had comparable behavioural recovery after injury. (**E-H**) Serial multifiber electrophysiology after sciatic nerve crush. Recovery of SNAP and CMAP amplitudes and conduction velocities were similar between the groups. (**I-N**) Morphological semithin section analysis of nerves and myelinated axons at baseline and 28 days follow a sciatic crush injury. Samples were analyzed distal to the crush zone. Baseline metrics, as above, show higher numbers of sural areas, sciatic and sural myelinated axon numbers. At 28 d, there were greater numbers of sciatic axons in sensory neuron PTEN-/-mice compared to wild type littermates. [(I) *p=0.018; (J)**p=0.004; (K) *p=0.02, *p=0.026;(L)**p=0.007 two-tailed Student’s t-tests; n=6 wild type, 5 PTEN-/-; (I) #p=0.04, (J) #p=0.026; (N)#p=0.05 one-tailed Student’s t-tests, n=6 wild type, 5 PTEN-/-]

At 28 days harvested sciatic and sural axons had larger nerve areas in PTEN-/-than littermate controls (**Figures 6I,J**). In the sciatic nerve regenerating myelinated axons were present with greater numbers of myelinated axons distal to the crush zone at the 28 day endpoint (**Figure 6K,L**). In sural axons there was a nonsignificant trend toward more axons at endpoint. Sciatic and sural nerve axon densities at 28 days following injury were similar between PTEN-/-mice and their littermate controls (**Figure 6M,N**).

Overall, these regeneration studies identified only modest impacts of PTEN deletion on recovery of thermal sensation at an early time point only and repopulation of sciatic axons. However other behavioural tests and electrophysiological indices after sciatic crush injury were comparable in the groups. While an impact on sensory axons alone might be anticipated, it may be that regenerative plasticity, already heightened prior to injury, was not further enhanced with added injury.

## Methods

### PTENflox-AdvillinCre mice and behavioural testing

Homozygous PTEN-flox (+/+) mice were bred with PTEN-flox+/-;AdvillinCre +/- or +/+ mice and screened for the PTEN-flox+/+;AdvillinCre+/-genotype in order to generate a conditional knockout of PTEN in adult sensory neurons. Male Advillin-Cre mice (Advillin^Cre/+^; Jackson Labs - #032536) were crossed with female Pten^Flox/Flox^ mice (Jackson Labs - #00644). Males of the preferred genotype (Advillin^Cre/+^:: Pten^Flox/+^) were selected from the progeny to breed in a second round with additional female Pten^Flox/Flox^ stock. Males with the Advillin^Cre/+^:: Pten^Flox/Flox^ genotype were subsequently used in this study. The Advillin^Cre/+^:: Pten^Flox/Flox^ males are viable and fertile and were therefore used for colony maintenance by breeding with Pten^Flox/Flox^ females to maximize the output. The genetic status of all mice was determined using a PCR reaction to detect either Advillin-Cre or Flox-Pten alleles as described in detail on the Jackson Labs website (Jax.org). In brief, Advillin-cre alleles were detected using the following primer sets: Advillin MF-5’-CCCTGTTCACTGTGAGTAGG-3’; Advillin MR-5’-AGTATCTGGTAGGTGCTTCCAG-3’; Cre MR - 5’-GCGATCCCTGAACATGTCCATC-3’; producing roughly 500 and 180 bp bands for wildtype and transgenic, respectively. Pten flox status was determined using a PCR reaction designed according to Jax.org. In brief, the following primers were used to detected flox-pten: oIMR9554 - CAA GCA CTC TGC GAA CTG AG; oIMR9555 - AAG TTT TTG AAG GCA AGA TCG; and yielded DNA bands at ∼156 and 338bp for wildtype and transgenic, respectively. AdvillinCre littermates were used as controls.

Mechanosensation was assessed using a Dynamic Plantar Aesthesiometer (Ugo Basile). Mice were placed on a metal grate, and a wire filament was pushed against the plantar surface of the hindpaw (ipsilateral to injury) with increasing force until paw withdrawal. Force and latency to withdrawal are recorded at the instant of paw withdrawal. The mean of 3 consecutive measures was used.

Thermal sensitivity was assessed using a Plantar Test (Hargreaves Apparatus) (Ugo Basile). Mice were placed on a transparent surface and an infrared heat source of increasing intensity is applied to the plantar surface of the hindpaw (ipsilateral to injury) until paw withdrawal. Latency was recorded at the instant of paw withdrawal. The mean of 3 consecutive measures was used.

Hindlimb grip strength was assessed using a Grip-Strength Meter (Ugo Basile). Mice were held by the tail while being allowed to grab onto plastic cup with their forelimbs. One hindlimb was allowed to grasp the bar, and the mouse is then pulled backwards until grip is released. The maximum force at the moment of grip release was recorded. The mean of 3 consecutive measures was used.

Gait was analyzed using the Digigait (Mouse Specifics, Framingham, MA) system. Mice are placed onto a transparent treadmill while their paw images are being recorded by a camera underneath. The mice were run on the treadmill at a speed of (15 cm/s) and recorded for 8-12 seconds. Gait analysis was performed using Digigait 12.4 software, reporting the sciatic functional index (SFI), which is a measure of gait impairment calculated from measurements of paw length, paw width, and toe spread. The protocols were reviewed and approved by local animal care committees following recommendations of the Canadian Council on Animal Care (CCAC).

### Nerve histology

Nerve tissue was fixed in 2.5% glutaraldehyde/0.025 M cacodylate buffer at 4°C overnight, then washed 6 times for 5 minutes each in 0.15 M cacodylate buffer. Fixed nerves were stained in a 2% Osmium Tetroxide/0.12 M cacodylate solution for 2.5 hrs at room temperature (RT), then washed 6 times for 5 minutes each in 0.15 M cacodylate buffer. Nerves were dehydrated using 5 ethanol washes of increasing concentration (70%, 80%, 95%, 100%,100%) for 8 minutes each, and then twice with 100% propylene oxide for 8 minutes each. The dehydrated nerves were then infiltrated overnight with a 50:50 (v:v) mixture of JEMBED:propylene oxide. The nerve tissue was embedded in JEMBED the following day and baked overnight at 45°C, and overnight again the next day at 65°C. Semithin sections were stained with toluidine blue and analyzed for cross-sectional area, axon profile count, axon density and size distribution and using ImageJ.

### Neuron cell culture, Immunocytochemistry

Primary sensory neuron cultures were prepared from L4–L6 dorsal root ganglia (DRGs) of C57BL/6 mice, including PTEN knockout and littermate control mice, to evaluate neurite outgrowth in vitro. DRGs were dissected and cultured in DMEM/F12 medium supplemented with appropriate growth factors. Neurons were seeded into 4-well chambered slides and allowed to adhere for 1 hour before being cultured for 24 hours under standard conditions. At 24 hours, cultures were fixed with 2% paraformaldehyde (PFA) for 10 minutes and processed for immunocytochemistry. Neurons were stained (after blocking with 1% BSA for 20 min) using a mouse monoclonal anti-β3-tubulin antibody (1:500; Sigma, Cat# T8578), followed by incubation with a goat anti-mouse secondary antibody conjugated to Alexa Fluor 488 (1:200; Invitrogen, USA, Cat# A-11001).Fluorescent images were acquired using an Axioskop fluorescence microscope (Carl Zeiss, Germany) at 20× magnification. Quantitative analysis of neurite outgrowth and additional regenerative parameters was performed using WIS-NeuroMath (http://www.cs.weizmann.ac.il/~vision/NeuroMath/index.html) software.

### Immunohistochemistry

Lumbar 4–6 dorsal root ganglia (DRGs) or sciatic nerves were immersed in Zamboni’s fixative at 4 °C overnight, followed by washing in phosphate-buffered saline (PBS). Tissues were then cryoprotected in 20% phosphate-buffered sucrose for 12 hours and embedded in Tissue-Tek OCT compound. Cryosections were cut at 12 μm thickness and processed for immunostaining. Sections were washed in PBS containing 0.05% Tween 20 (PBS-T) and blocked in a solution of 1% bovine serum albumin (BSA) and 5% goat serum for 30 minutes at room temperature. Primary antibodies were applied in a humidified chamber and incubated for 2 hours at room temperature. Following primary antibody incubation, sections were incubated with the following secondary antibodies for 90 minutes at room temperature: Alexa Fluor 488 goat anti-mouse IgG (1:200; Invitrogen, USA) or Alexa Fluor 546 goat anti-rabbit IgG (1:200; Invitrogen, USA). After washing, sections were mounted using VectaShield with DAPI (Vector Laboratories, USA) to visualize cell nuclei. Primary antibodies included: (i) PTEN: Rabbit polyclonal anti-PTEN (1:500; Santa Cruz Biotechnology, Cat# sc-7974) - Colocalized with NF200 to calculate integrated fluorescence; (ii) NF200: Mouse monoclonal anti-NF200 (1:600; Sigma, Cat# N4142); (iii) IB4: (1:500; [Cat# Sigma-Aldrich L2140]) - Colocalized with β3-Tubulin - Calculated % IB4 positive cells counted using Image J; (iv) β3-Tubulin: Mouse monoclonal anti-β3 tubulin (1:300; Sigma, Cat# T8328); (v) ATF3: Rabbit antibody (1:300; Abcam, Cat# ab207434); (iv) Phospho-RAC: Mouse antibody (1:300; (1:300; [Cat# 26903])-Colocalized with PGP9.5; (v) PGP9.5: Rabbit anti-PGP9.5 (EnCor Biotech Inc., Gainesville, FL; [Cat# RPCA-UCHL]); SCG10: Rabbit antibody (1:500; Invitrogen, Cat# 720178); MBP: (1:300; ThermoFisher [Cat# 110008])-colocalized with PGP on transverse sciatic nerve sections-counted 100 cells in sections of sciatic nerves and represented the respective expression as %. Using ImageJ software, five images per animal (n = 4) were analyzed. For each image, the mean gray value was measured to determine integrated fluorescence intensity per area, and positive cells were counted.

### Electrophysiology

Compound muscle action potentials (CMAPs) and sensory nerve action potentials (SNAPs) were recorded in anaesthetized mice. During all recordings, mice were under 2% isoflurane gas anaesthesia and a heat lamp to maintain a near nerve temperature of 37.0±1.0^0^ C. A grounding electrode was inserted at the upper thigh with all stimulations. For CMAPs, positive and negativere cording electrodes were placed in the foot with the positive placed 2-3 mm distal to the negative. Stimulation occurred separately at two sites: sciatic notch, and the knee. The reference stimulating electrode was inserted 2-3 mm proximal to the negative. A current was used to supramaximally stimulate the sciatic nerve at each location and the distance between the active stimulating electrodes at each site was recorded. Latency, amplitudes and conduction velocity from each recording were calculated. SNAPs were acquired with stimulation at the foot (positive more distal) and recording at the knee (positive more proximal), and the distance between the two negative electrodes were recorded. The average of 20 SNAP recordings were used to control for background noise.

### Crush injury model

Mice underwent a unilateral sciatic nerve crush under 2% isoflurane gas anaesthesia. A 5 mm incision was be made in the skin at mid-thigh level, and the sciatic nerve exposed by blunt dissection through the muscle. The nerve was then crushed for 20s using fine-tipped needle drivers. Incisions were closed with 4-0 vicryl sutures. Post-operative mice received buprenorphine jelly (5 mg/kg) for 3 days (day 0, 1, 2). At 28 days, mice were sacrificed by cardiac puncture. The following tissues (both contralateral and ipslateral to injury) were harvested for histological analysis: sciatic nerve, sural nerve, DRGs (L4, 5, 6), hindpaw footpad.

### Statistical Analysis

Results were reported as means±SEM. Differences in measures were assessed using Student’s t-test (one or two tailed as appropriate). Regeneration measures were compared using repeated measures, two-way ANOVA. Statistical significance was taken at p < 0.05.

## Discussion

One of the knowledge gaps in neurodevelopment and regeneration is how molecular control of ongoing growth is restrained or shut down during full adulthood. Ongoing remodeling in adult axon innervation would be necessary to compensate for axon loss from minor injuries, skin or endothelial cell turnover. However, the major assumption is that the nervous system including brain, spinal cord and peripheral axons retain stability to ensure reliable function. In advanced age, these connections may decline. In the case of peripheral axons, their coordinated signaling during activities of daily living require a uniformity of structure and function. Moreover use of this sensation in retaining memories in the CNS requires consistent input. Here we report that deletion of a single signaling molecule, namely PTEN, disrupts this uniform stability in how sensory axons are restrained within fascicles.

The relationship between PTEN and regenerative plasticity has now been well established for a number of years, originally in both optic nerve regrowth and in our own work in peripheral nerves ^7,21^. PTEN is hypothesized to ‘brake’ regenerative regrowth by disrupting the connection between PI3K and pAkt signaling, a central mechanism of cellular survival and growth signaling. It is likely that many, if not most growth factor actions operate through pathways that might be suppressed by PTEN. Moreover, there is a relationship, albeit of unclarified importance, between PTEN and the Myc interactome translational signaling network, specifically Mad1^23^. While not tested here, PTEN null neurons might exhibit Myc activation but the activation is complex since Mad1, an inhibitor of Myc also inhibits PTEN ^24^. A series of growth suppressive signals, largely studied in the oncology sphere, or tumour suppressors are nonetheless expressed in neurons. Their interruption enhances regenerative growth in peripheral neurons. Other examples are Mad1 suppressing Myc, but also Rb1 suppressing E2F1 and APC suppressing β-catenin ^5,8,23^.

In isolated adult neurons *in vitro*, there are additive benefits of growth cone modulation by activating the growth cone GTPase Rac1 combined with PTEN inhibition^1^. However PTEN expression at axon terminal tips and growth cones is less prominent. *In vivo*, applying local PTEN knockdown distally at the site of regrowing terminal axons into skin offered only modest impacts beyond those of a Rac1 activator. Moreover, in a growth cone turning assay, local application of a PTEN inhibitor to growth cones had negligible direct impacts on growth cones but application to the perikarya facilitated outgrowth in axons remote from the site of application ^9^. These findings indicate that PTEN suppression of growth may largely act within perikarya rather than at the growth cone. Activation of outgrowth likely involves a disseminated intracellular signal through pAkt which is identifiable in growth cones and activates Rac1.

What is remarkable in the current work is the impact of selective sensory neuron PTEN deletion on distal axon behaviour, without an obvious DRG or perikaryal structural change beyond ATF3 nuclear expression. The evidence for inappropriate but robust axon sprouting in sensory neuron PTEN-/-mice in this work was supported by several findings. *In vitro* worked confirmed that sensory neurons lacking PTEN had heightened outgrowth, resembling findings from previous work in wild type neurons with PTEN inhibition or knockdown ^7^. In mice lacking sensory neuron PTEN, peripheral nerve trunks were enlarged with greater numbers of myelinated axons forming clusters. Sprouting was not uncontrolled, as these events were confined to endoneurial fascicles without aberrant growth into the perineurium. Uncontrolled axon growth into the epineurium is frequently observed in neuromas and other nerve injuries ^32,33^. Numbers of myelinated axons were increased in the sciatic and sural nerves. The clusters identified sprouting likely from both large caliber and small caliber neurons. Electrophysiological studies also confirmed myelinated axon expansion giving rise to an increased amplitudes of SNAPs in PTEN-/-sensory mice. The amplitudes of SNAPs are an electrophysiological index of viable sensory axons at the site of the recording electrode. These findings indicated that the new axons were electrically excitable, as expected from normal maturation. ATF3 nuclear expression, a marker of regenerative growth was not confined to small neurons. Similarly, myelinated axon sprouts would not be expected to arise from parent unmyelinated axons of small caliber neurons. IB4 is of particular interest because of its particularly high expression of PTEN in wild type sensory neurons. However, previous work indicated PTEN is widely expressed beyond this nonpeptidergic population.

Ultrastructural studies added several key findings. Firstly, they confirmed the unexpected clustering of atypical endoneurial axons. Secondly they illustrated that apparent sprouts were both small nascently myelinated axons, smaller than A delta axons, but typical of early maturing regenerative sprouts. In original ultrastructure studies of regenerating nerve, Morris and colleagues ^16^ distinguished morphological criteria of regenerating units of axons and Schwann cells (SCs) growing from the proximal stumps of transected nerves. A mature appearing myelinated axon that shares its SC and basement membrane with one or several unmyelinated axons was classified as Type I regenerative unit, with axons closely apposed to the SC membrane. Clusters of unmyelinated axons alone were classified as Type II regenerative units. The latter however differed from Remak unmyelinated axon bundles of normal nerves, lacking leaves of SC cytoplasm between axons in normal nerves. In AdvCre;PTEN-/-mice axons within these units were of varying caliber, closely apposed and numbered from a single axon up to several. Most of the units identified ultrastructually in our work would be classified as Type II units with only rare examples of Type I units. The caveat to this observation however was that Type I units here were only seen accompanying small myelinated axons, rather than mature myelinated axons Clusters of axons labeled by PGP9.5 included both small myelinated axons, indicated by the coexpresion of MBP but also profiles devoid of MBP sheaths also indicating the presence of unmyelinated axons. Finally, increased numbers of unmyelinated axons innervating the epidermis were encountered. Taken together, the ultrastructural work and immunohistochemical studies confirmed that axon expansion included clusters of both myelinated and unmyelinated axons. Finally we also identified axons expressing SCG10 a marker of sensory axon regenerating fibers, not expected of intact nerves ^14,25^.

Despite the rises in numbers of sensory axons, mice with sensory neuron PTEN deletion had intact mechanical and thermal sensation without a change in gait. Motor grip power and motor electrophysiology were normal, in keeping with the conditional deletion only in peripheral sensory neurons. At first glance, one might expect hypersensitivity to thermal sensation related to unmyelinated fiber expansion or to mechanical sensitivity given the rises in myelinated axons. However, supernumerary sprouts from individual parent sensory neurons and axons may not convey additive sensory information. While axon sprouting has been associated with pain phenotypes, this has largely been in the setting of injury and regenerative or collateral sprouting. Once adult circuitry is established there may be little benefit from added axons. In the epidermis, there are constraints to axon numbers per unit of epidermal length and addition may not increase sensation.

Parallel expansions in neural structures has been described in the CNS. Kazdoba and colleagues ^10^ described massive enlargement of the forebrain and neonatal mortality in mice undergoing deletion of PTEN specifically in early postmitotic neurons in the developing forebrain. These findings differed however from the current work in which plasticity was expressed in axons of stable adult mice without rises in numbers of ganglion neurons. We had evidence of ongoing growth, as apposed to a stable rise in developmental expression. Kwon et al ^11^ reported findings in mice with a neuron-specific enolase (Nse) Cre recombinase to generate CNS PTEN deleted mice. These mice survived and exhibited macrocephaly, hippocampal neuronal hypertrophy, with hypertrophic and ectopic dendrites and heightened axonal synapse formation. Mossy fiber tracts were enlarged. These changes accompanied hyperactivity in stressful conditions, anxiety-like behaviour, impaired learning and seizures. Evidence of ongoing hippocampal axonal growth were akin to our observations in adult peripheral neurons. In the CNS,PTEN has been shown to associate with the pro-apoptotic protein Bax, promoting its entry to mitochondria where it initiates apoptosis. Knockdown of PTEN inhibits apoptosis ^31,27^. Bax activity, in turn, has been shown to be required for developmental apoptosis in DRG neurons during embryonic days E11.5 - E14.5. Further, Bax knockout mice exhibit increases in the number of myelinated axons in peripheral (facial) nerve and displayed a pattern of smaller-diameter myelinated axons that resembled the axon distributions observed in our work ^29^. Finally, in motor neurons conditional PTEN deletion using a Chat Cre recombinase generated hypertrophy of motor neurons, axons and nerves, in a parallel fashion to that presented here. However, we did not demonstrate sensory neuron hypertrophy or larger caliber axons, but instead evidence of ongoing regenerative sprouting. As in the case of PTEN pharmacological inhibition or siRNA, the motor PTEN-/-mice demonstrated improved axon regeneration, but limited to motor neurons, specifically within the facial nerve.

Unlike the impact of exogenous PTEN knockdown or inhibition on enhancing regeneration *in viv*o, mice with sensory neuron PTEN deletion displayed only modest enhancement of sensory recovery. Motor recovery, as discussed, was not expected. We did note a borderline and transient improvement in thermal sensation at 14 days in the PTEN-/-mice not sustained to 28 days. In addition, sciatic axon numbers were higher in the regenerating nerves at endpoint after injury. Thus, there were some features of heightened axon regrowth in the PTEN null mice but less striking than observed in other PTEN work. PTEN deletion did not convey any definite and persistent benefits in mechanical or thermal sensation. One possibility is that PTEN-/-sensory neurons are already possessed of a regenerative activated state. Given this, there may be no additional benefits from added nerve injury available to drive enhanced growth and RAGs.

**Supplemental Figure 1:**
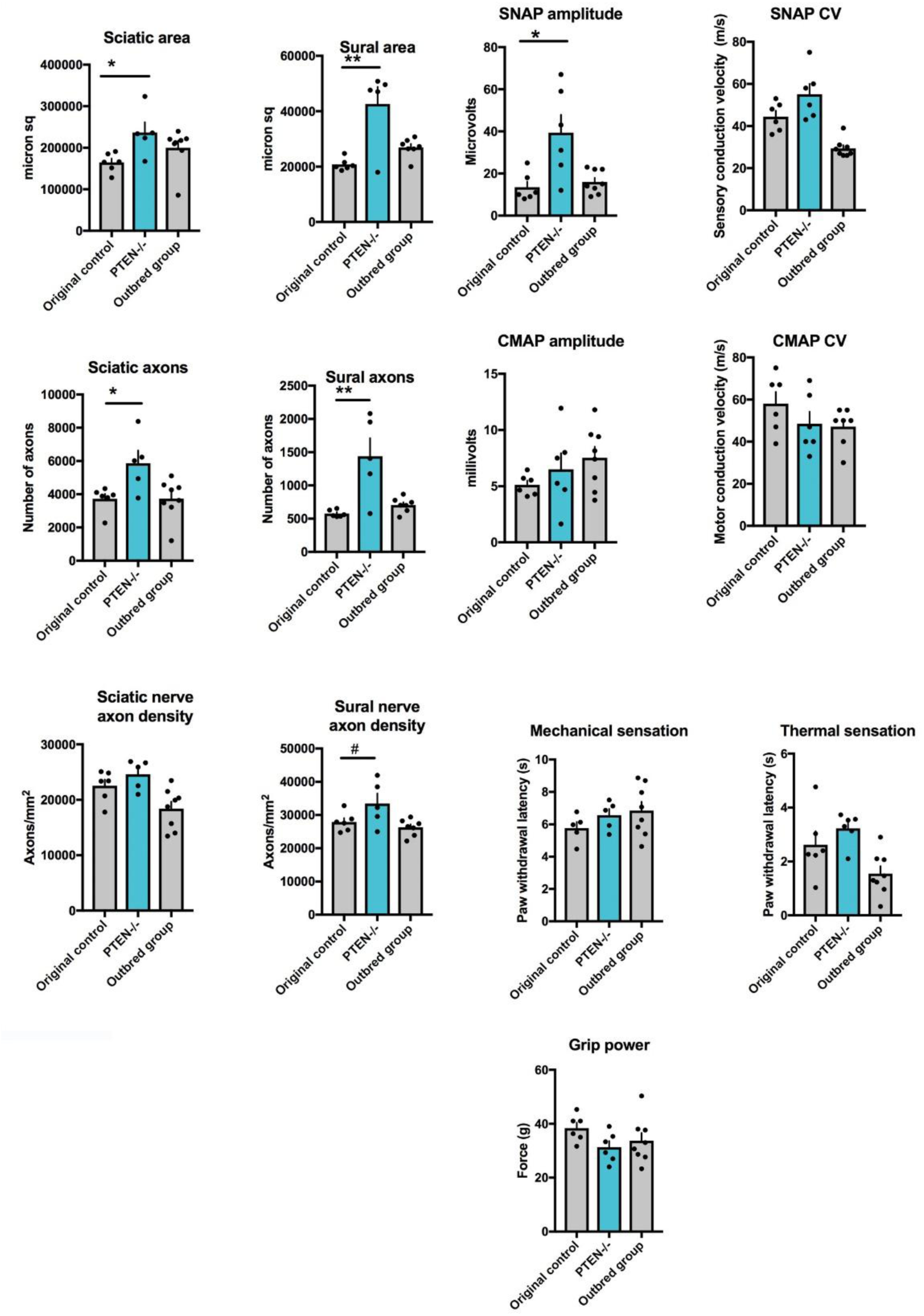
Original cohort of sensory PTEN-/-mice compared to an additional control group from this cohort line with subsequent loss of the genotype. Retention of PTEN expression in this cohort was confirmed by immunohistochemistry (not shown). As in the comparisons to concurrent controls in Figure 2 and 5, sensory PTEN-/-mice had larger areas of sciatic and sural nerves, greater numbers of axons, rises in SNAP amplitudes but no significant changes in sensory or motor conduction velocities, CMAP amplitudes or behavioural measures of sensation and grip power compared to these loss of genotype controls.

